# Amino acid competition shapes *Acinetobacter baumannii* gut carriage

**DOI:** 10.1101/2024.10.19.619093

**Authors:** Xiaomei Ren, R. Mason Clark, Dziedzom A. Bansah, Elizabeth N. Varner, Connor R. Tiffany, Kanchan Jaswal, John H. Geary, Olivia A. Todd, Jonathan D. Winkelman, Elliot S. Friedman, Babette S. Zemel, Gary D. Wu, Joseph P. Zackular, William H. DePas, Judith Behnsen, Lauren D. Palmer

## Abstract

Antimicrobial resistance is an urgent threat to human health. Asymptomatic colonization is often critical for persistence of antimicrobial-resistant pathogens. Gut colonization by the antimicrobial-resistant priority pathogen *Acinetobacter baumannii* is associated with increased risk of clinical infection. Ecological factors shaping *A. baumannii* gut colonization remain unclear. Here we show that *A. baumannii* and other pathogenic *Acinetobacter* evolved to utilize the amino acid ornithine, a non-preferred carbon source. *A. baumannii* utilizes ornithine to compete with the resident microbiota and persist in the gut in mice. Supplemental dietary ornithine promotes long-term fecal shedding of *A. baumannii*. By contrast, supplementation of a preferred carbon source—monosodium glutamate (MSG)— abolishes the requirement for *A. baumannii* ornithine catabolism. Additionally, we report evidence for diet promoting *A. baumannii* gut carriage in humans. Together, these results highlight that evolution of ornithine catabolism allows *A. baumannii* to compete with the microbiota in the gut, a reservoir for pathogen spread.

## Introduction

Antimicrobial resistance (AMR) is one of the greatest threats to human health worldwide. AMR was associated with approximately 5 million deaths in 2021.^1^ Pathogens encoding AMR often persist in hospitals and other healthcare settings by asymptomatically colonizing the microbiota of patients and staff.^2,3^ *A. baumannii* is a healthcare-associated opportunistic pathogen and was the second leading cause of deaths directly attributable to AMR in 2021.^1,4^ In 2024, the World Health Organization again listed carbapenem-resistant *A. baumannii* as the highest-priority target for antibiotic research and development due to increased mortality rates.^5^ *A. baumannii* can colonize any site in the body and asymptomatic colonization is associated with increased risk of infection and overall mortality in hospitalized patients.^4,6–11^ As early as 1993, the gut was identified as a reservoir for *A. baumannii* in intensive care unit patients, as gut colonization preceded nasopharyngeal colonization and clinical infection.^12^ While *Acinetobacter* spp. gut carriage is low prevalence in non-hospitalized populations (<1% of people),^13^ gut colonization with *A. baumannii* can be detected in up to 41% of hospitalized patients.^8,10,12,14–18^

Gut colonization increases the risk of clinical infection and transmission. Prospective culture-based studies showed that fecal colonization with multidrug resistant (MDR) *A. baumannii* increased the risk of clinical infection 5.2-15.2 fold.^6,9,10^ *A. baumannii* strains that asymptomatically colonize the gut have been isolated from clinical infections in the same patients or after transmission to other patients.^7,19–24^ Moreover, rectal swabs detected carbapenem-resistant *A. baumannii* at a higher rate than axillary or nasal swabs in asymptomatically colonized patients,^16^ emphasizing the gut as a potential reservoir for antimicrobial-resistant strains. In addition to colonizing adults in healthcare settings, *A. baumannii* has been isolated from the neonatal gut and is a threat to infant health.^17,25–27^ In summary, *A. baumannii* colonizes the gut asymptomatically in healthcare settings, which increases the risk of clinical infections.

Despite evidence for the role of *A. baumannii* gut colonization in patients, few mechanistic studies have explored this phenomenon and the *A. baumannii* niche in the gut ecosystem is unclear. Previous studies have demonstrated persistent carriage in mouse models with antibiotics and identified a role for secretory IgA and *A. baumannii* thioredoxin in short-term carriage.^28–31^ However, little is known about the strategies that *A. baumannii* employs to colonize the gut.

For *A. baumannii* to colonize the gut, it must overcome colonization resistance from the resident microbiota, which is often mediated by niche exclusion via competition for nutrients.^32–34^ Thus, *A. baumannii* metabolic processes are likely important determinants for colonization and persistence in the gut.^35–37^ Here, we show that ornithine catabolism contributes to *A*. *baumannii* gut colonization in a post-antibiotics (post-abx) mouse model. We uncovered a second arginine succinyl transferase pathway (AST) operon present only in pathogenic *Acinetobacter* species that encodes a predicted ornithine succinyltransferase (AstO) necessary for catabolizing ornithine. In a post-abx mouse model, *A. baumannii* AstO-dependent ornithine catabolism is required to compete with the resident microbiota to persistently colonize the gut. We report that dietary supplementation of ornithine or monosodium glutamate, a preferred carbon source, promotes *A. baumannii* colonization in mice and an association between diet and *A. baumannii* abundance in the gut microbiota of human infants. These findings highlight a metabolic strategy that allows *A. baumannii* to utilize a non-preferred carbon source in the gut environment, which is thought to be a reservoir for pathogen spread in healthcare settings.

## Results

### A partial second AST pathway operon confers ornithine catabolism and is critical for *A. baumannii* gut colonization

The molecular mechanisms that govern how *A. baumannii* colonizes the gut are largely unknown. Because competition for nutrients is central to pathogen colonization of the gut, we predicted that *A. baumannii* and related pathogenic *Acinetobacter* spp. may have evolved additional metabolic capacities to colonize the gut. We identified a partial second AST pathway operon (*astNOP)* in pathogenic *Acinetobacter* species that is absent in non-pathogenic *Acinetobacter* species (Figure 1A and S1A). The AST pathway is typically encoded in a single polycistronic operon *astCADBE* and degrades L-arginine to L-glutamate in *Escherichia coli, Pseudomonas aeruginosa,* and *Klebsiella aerogenes*.^38–40^ In *A. baumannii* ATCC 17978, the second *ast* locus encodes homologs of AstC (Ast<u>N</u> for <u>N</u>_2_-succinylornithine transaminase), AstA (Ast<u>O</u> for <u>o</u>rnithine succinyltransferase), a putative amino acid permease (Ast<u>P</u> for predicted <u>p</u>ermease) and a divergently transcribed predicted Lrp/AsnC-family regulator (Ast<u>R</u> for predicted regulator) (Figure 1A and S1A). In *E. coli* and *P. aeruginosa*, AstA can utilize arginine and ornithine as a substrate to enter the AST pathway.^38,39^ Therefore, we hypothesized that AstO is an ornithine succinyltransferase that allows *A. baumannii* to utilize ornithine (Figure 1B). To test this, we constructed *A. baumannii* ATCC 17978 *astA* and *astO* mutant strains. Multiple attempts to delete *astA* led to polar effects on downstream genes in the operon; therefore, an *astA*^L125A,H229A^ mutant (*astA* mutant) was constructed with substitutions at predicted catalytic residues.^41^ WT grew with either arginine or ornithine as the sole carbon source (Figure 1C). The *astA* mutant grew with ornithine but not arginine as the sole carbon source (Figure 1C). The Δ*astO*::Kn (Δ*astO*) mutant grew with arginine but did not grow with ornithine as the sole carbon source (Figure 1C). The Δ*astO* mutant defect in growth with ornithine was complemented by *astO* expression *in trans* (Figure S1B). These data suggest *A. baumannii* uses AstA to catabolize arginine and AstO to catabolize ornithine. The double mutant *astA* Δ*astO* was not able to utilize either arginine or ornithine (Figure 1C), demonstrating that under these conditions the AST pathway is the only arginine/ornithine catabolic pathway in *A. baumannii*.

**Figure 1.**
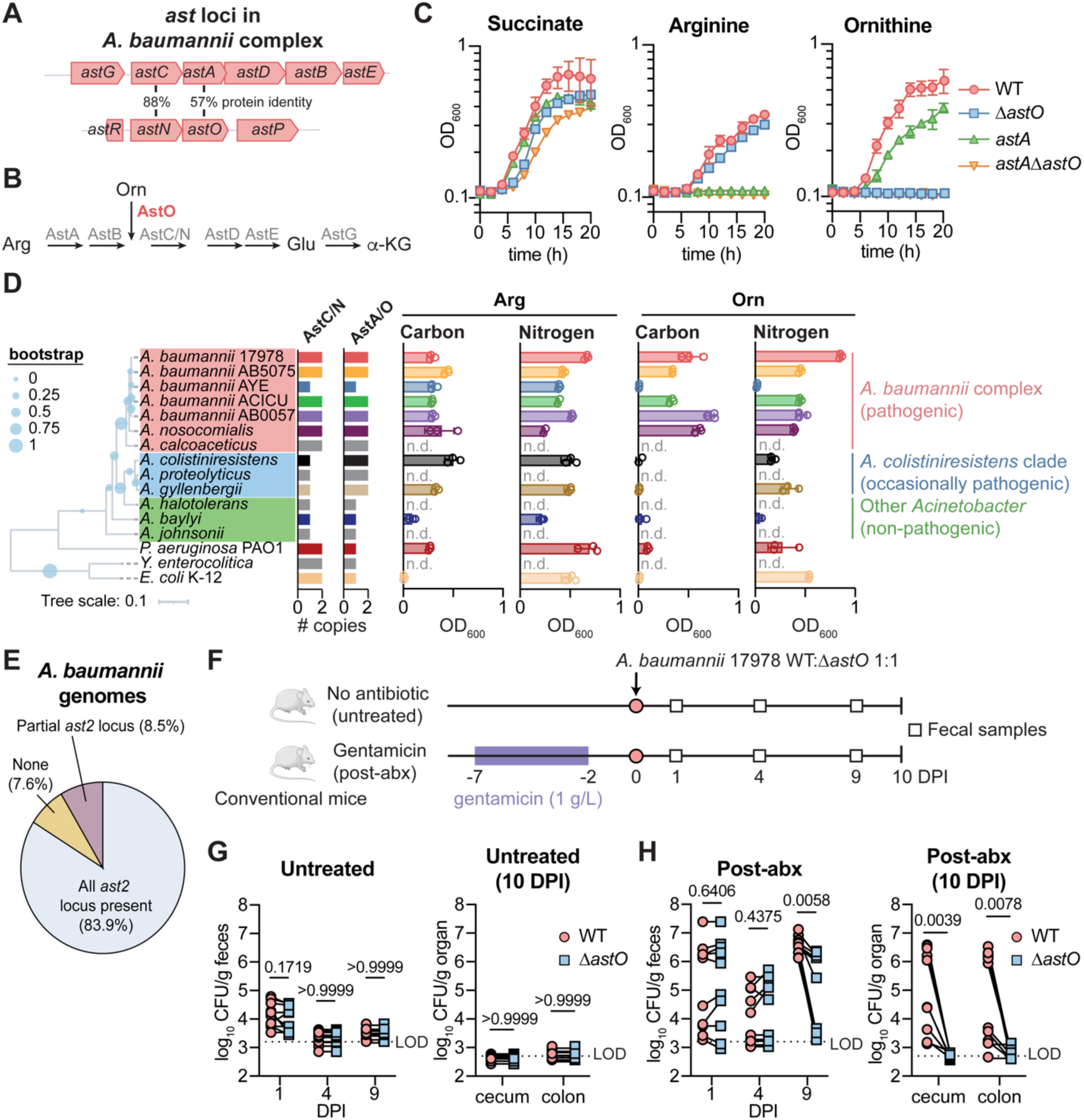
A partial second AST pathway operon confers ornithine catabolism in pathogenic *Acinetobacter* species and contributes to *A. baumannii* gut colonization. (A) *ast* loci in *A. baumannii* complex (ABC). (B) Schematic of the predicted arginine succinyl transferase (AST) pathway in *A. baumannii*. (C) *A. baumannii* 17978 WT, Δ*astO*, *astA*^L125A^ ^H229A^ and *astA*^L125A^ ^H229A^ Δ*astO* strains were grown in M9 minimal media with arginine, ornithine, or succinate as the sole carbon source. Growth was monitored by optical density at 600 nm (OD_600_) (n = 3, mean ± SD, experiments were repeated at least 6 times with similar results). (D) Phylogenetic species tree with inferred copy numbers of AstA/O and AstC/N (tree scale in amino acid substitutions). Max OD_600_ of cultures grown with arginine or ornithine as the sole carbon or nitrogen source in M9 medium with succinate (glucose for *E. coli*) or NH_4_ as controls. Growth was over 20 h (n = 3, mean ± SD, experiments were repeated at least 2 times with similar results). (E) Prevalence of the *ast2* operon in genomes of *A. baumannii* isolates. (F) Experimental design of post-abx *A. baumannii* 17978 gut colonization model. (G-H) *A. baumannii* 17978 CFU from fecal samples at 1, 4, and 9 DPI and in the cecum and colon at 10 DPI (n = 10 female Swiss Webster mice combined from 2 independent experiments; *p* by Wilcoxon test with Holm-Sidak’s multiple comparisons). Arg, arginine; Orn, ornithine; Glu, glutamate; α-KG, α-ketoglutarate; n.d., not determined; DPI, days post inoculation; LOD, average limit of detection.

We next investigated the phylogeny and conservation of *astO* and ornithine catabolism among the *Acinetobacter* genus and *A. baumannii* isolates. The *astCADBE* operon is ubiquitously encoded in the *Acinetobacter* genus. The *Acinetobacter baumannii* complex includes all significant pathogens and encodes *astNOP* with divergent *astR* locus (red, Figure 1A, D and S1A). An additional group that includes occasional pathogens encodes only *astO* near an unrelated LysR-type regulator (*A. colistiniresistens* clade, blue, Figure 1D and S1A).^42^ *A. colistiniresistens astO* can partially complement the Δ*astO* defect in growth on ornithine as the sole carbon source, demonstrating it is an ornithine succinyltransferase (Figure S1B). Other clades of *Acinetobacter* that are non-pathogenic encode only the *astGCADBE* operon (Other *Acinetobacter*, green, Figure 1D and S1A). Because the AstO activity in pathogenic *Acinetobacter* clades is similar to activity present in other bacteria such as *E. coli* and *P. aeruginosa*, we considered the hypothesis that an ancestor of *A. baumannii* acquired the *astO* from another organism via horizontal gene transfer (HGT). Phylogenetic analysis showed that AstA and AstO proteins in the pathogenic ABC clade and *A. colistiniresistens* clades were more closely related to each other than to AstA encoded by *Yersinia enterolitica, P. aeruginosa*, or *E. coli* (Figure S1C). This suggests that *A. baumannii astO* was acquired via HGT from a closely related species or from a partial duplication after the divergence of pathogenic *Acinetobacter* clades.

To test if AstO was associated with the ability to use ornithine as the sole carbon and/or nitrogen source in pathogenic *Acinetobacter* species, we quantified AstC/N and AstA/O homologs among select *A. baumannii* strains and the reference genomes of *Acinetobacter* species. Encoding only one homolog indicated the presence of only the *astGCADBE* operon, while encoding two homologs indicated presence of a second *ast* locus (Figure 1D). We next measured whether selected strains and species could use arginine and/or ornithine as the sole carbon and energy source in minimal media. Encoding AstO (*i.e.*, encoding two homologs of AstA/O) correlated to whether the *Acinetobacter* species could use ornithine as the sole carbon and/or nitrogen source (Figure 1D). Together, these data demonstrate an association between encoding AstO and ornithine utilization and suggest that the *ast2* locus is associated with pathogenicity in *Acinetobacter* species.

To determine whether second *ast* locus is conserved among *A. baumannii* isolates, the prevalence of the *astR* and *astNOP* was assessed among our previously described set of 229 *A. baumannii* genomes deduplicated to remove clonal lineages (*e.g.* outbreak sequencing).^43^ The complete *ast2* locus was present in 83.9% of *A. baumannii* isolates, while 8.5% of *A. baumannii* isolates encoded a partial *ast2* locus, and 7.6% of *A. baumannii* isolates lack the *ast2* locus (Figure 1E, S1D). Therefore, the *ast2* locus is conserved in the majority of sequenced *A. baumannii* isolates, suggesting AstO-dependent ornithine catabolism may confer a fitness advantage to *A. baumannii*.

Ornithine has emerged as key nutrient for gut colonization by the antimicrobial-resistant pathogen *C. difficile*.^44–47^ To test whether *A. baumannii* uses ornithine catabolism for gut colonization, we developed a post-antibiotics (post-abx) mouse model of *A. baumannii* gut colonization due to the association between previous antibiotic treatment and *A. baumannii* gut colonization in humans.^9,20,23,48,49^ Female mice were administered 1 g/L gentamicin in the drinking water for 5 days or a vehicle treatment (untreated; Figure 1F). After 2 days recovery with normal drinking water, mice were orogastrically inoculated with 10^9^ CFU of a 1:1 mixture of the *A. baumannii* 17978 WT and Δ*astO* (Figure 1F).

Consistent with modeling asymptomatic colonization, there were no clinical signs of infection such as hunching or weight loss (Figure S1E). There was no significant difference in the burdens of *A. baumannii* between WT and Δ*astO* in the feces at 1 DPI in the post-abx group or untreated group (Figure 1G and 1H). *A. baumannii* was below the limit of detection (LOD) after day 4 in untreated mice, suggesting substantial colonization resistance by the intact resident microbiota (Figure 1G). However, there was *A. baumannii* gut colonization throughout the experiment in the post-abx group (Figure 1H). WT *A. baumannii* significantly outcompeted the *ΔastO* mutant by 9 DPI (Figure 1H). These data suggest that *A. baumannii* ornithine catabolism is required for persistent gut colonization in this post-abx model. At 10 DPI, WT *A. baumannii* CFU were significantly higher than the Δ*astO* mutant strain in the cecum and colon (Figure 1H). There was also a trend toward higher WT CFU compared to Δ*astO* in the small intestine at 10 DPI (Figure S1F). There were no *A. baumannii* CFU detected in the spleen at 10 DPI, suggesting *A. baumannii* did not disseminate to other organs (Figure S1G). Similar results were observed in males, demonstrating that this phenomenon was not sex-dependent (Figure S1H). We next tested whether *A. baumannii* AstO-dependent catabolism was also important for infection. However, Δ*astO* mutants had no defect in mouse models of lung infection or bloodstream infection (Figure S1I-J), the most common types of *A. baumannii* infections.^50,51^ Collectively, these findings suggest that the AstO- dependent ornithine catabolism is conserved in pathogenic *Acinetobacter* spp. and is important for *A. baumannii* persistence in the gut.

### Supplemental dietary ornithine promotes long-term *A. baumannii* gut colonization and fecal shedding in mice

Based on the ornithine requirement for persistent *A. baumannii* gut colonization, we hypothesized that supplementation of ornithine in the drinking water would promote long-term *A. baumannii* gut colonization. Two groups of female mice were orally inoculated with a 1:1 mixture of WT and Δ*astO A. baumannii* 17978. One group was supplemented with dietary ornithine HCl (1% w/v) in the drinking water 1 day prior to inoculation and both groups were monitored for 10 weeks (Figure 2A). This concentration was previously shown to promote asymptomatic *Clostridoides difficile* colonization.^45^ Without supplemental ornithine, *A. baumannii* WT outcompeted Δ*astO* from 10-42 DPI (Figure 2B).

**Figure 2.**
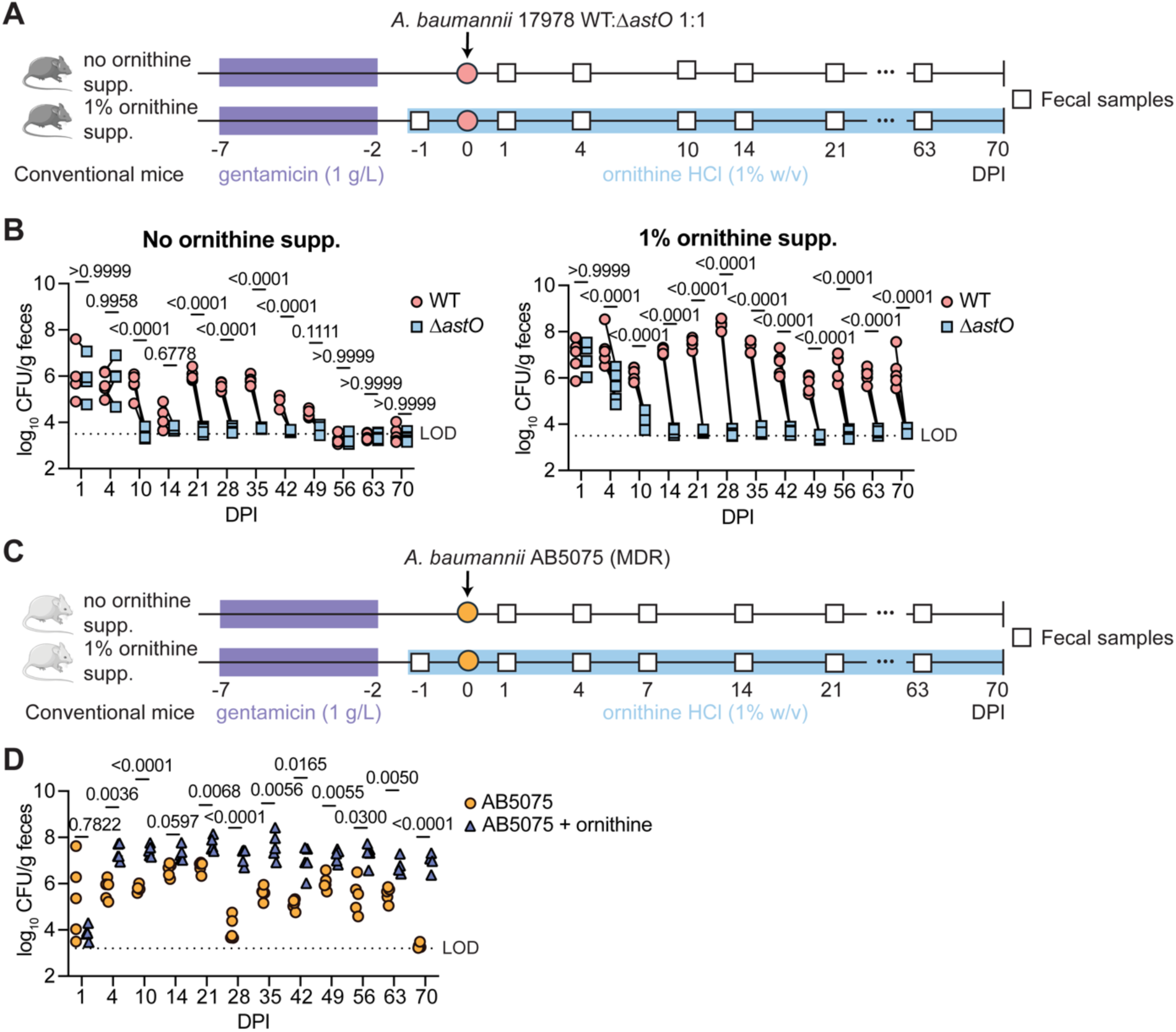
Supplemental dietary ornithine promotes long-term *A. baumannii* gut colonization and fecal shedding in mice. (A) Experimental design for ornithine supplementation and *A. baumannii* 17978 gut colonization. (B) *A. baumannii* 17978 CFU from fecal samples at the indicated DPI (n = 5 female C57BL/6J mice, *p* by two-way ANOVA with Sidak’s multiple comparisons; experiments were repeated 2 times with similar results). (C) Experimental design for ornithine supplementation and *A. baumannii* AB5075 inoculation. (D) *A. baumannii* AB5075 CFU from fecal samples the indicated DPI (n = 5 female Swiss Webster mice, *p* by two-way ANOVA with Sidak’s multiple comparisons). DPI, days post inoculation; LOD, average limit of detection.

Around 56 DPI, *A. baumannii* WT fecal shedding decreased to below the LOD (Figure 2B). By contrast, in mice given supplemental dietary ornithine in the drinking water *A. baumannii* 17978 WT colonized the gut to at least 70 DPI, while the Δ*astO* mutant was at or below the LOD from 14 DPI (Figure 2B and S2A). Together, these data suggest that *A. baumannii* AstO-dependent ornithine catabolism is required for long-term colonization and that dietary ornithine supplementation promotes persistent *A. baumannii* colonization. To determine whether ornithine supplementation similarly promotes persistent gut colonization by a recent MDR clinical isolate, we tested the effect of dietary ornithine supplementation on gut colonization by *A. baumannii* AB5075 (Figure 2C).^52^ The group receiving dietary supplementation of ornithine in the drinking water had significantly higher *A. baumannii* AB5075 gut colonization than the control group from 4 DPI (Figure 2D). *A. baumannii* AB5075 colonized to 63 DPI the gut in the control group and at least 70 DPI in the supplementary ornithine group (Figure 2D and S2B). Compared to *A. baumannii* 17878 inoculation, AB5075 colonized to higher levels and persisted longer in the gut (Figure 2B and 2D). Moreover, *A. baumannii* disseminated to the spleen at 70 DPI when mice inoculated with AB5075 were given supplementary ornithine (Figure S2B). This suggests that supplemental dietary ornithine and/or high bacterial burdens in the gut promotes *A. baumannii* AB5075 dissemination.

Together, these data demonstrate that supplemental dietary ornithine promotes long-term persistent gut colonization and may promote dissemination by *A. baumannii*.

### In the absence of the microbiota, there is no fitness advantage for AstO-dependent ornithine catabolism

Since *A. baumannii* colonization relied on microbiota disruption, we next explored whether *A. baumannii* may utilize ornithine to compete with the microbiota. First, the microbiota composition was profiled in fecal samples from the untreated and post-abx mice at 0 DPI and 9 DPI in Figure 1 by 16S rRNA gene sequencing. Microbiota α-diversity, a measure of richness and evenness on a local scale (*e.g.* in a single mouse), was reduced at 0 DPI in post-abx mice compared to untreated mice but was the same in post-abx and untreated mice by 9 DPI (Figure 3A). β-diversity, a measure of richness and evenness across samples, and relative abundance of amplicon sequence variants (ASV) were dramatically altered in the post-abx group compared with untreated group on day 0 and largely recovered by 9 DPI (Figure S3A- B). Together, these data demonstrate that gentamicin treatment dramatically alters the microbiota at the time of inoculation, but that the resident microbiota is largely restored by 9 DPI when *A. baumannii* required ornithine catabolism.

**Figure 3.**
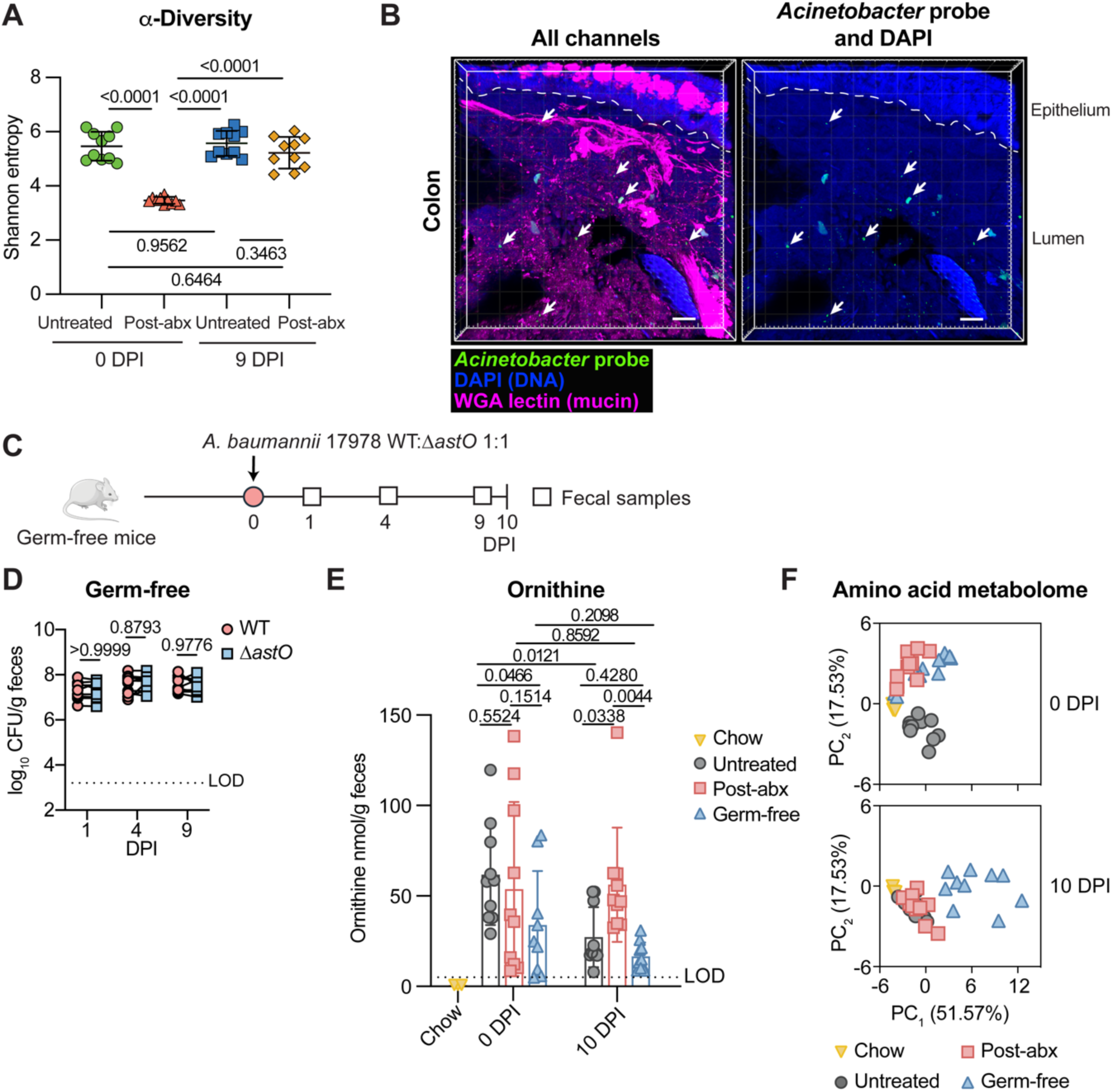
Microbiota diversity associates with *A. baumannii* AstO-dependent ornithine utilization and the gut amino acid metabolome. (A) α-diversity was calculated from 16S rRNA gene sequencing at 0 DPI and 9 DPI of *A. baumannii* 17978 WT and Δ*astO* (n = 10; data combined from mice in Figure 1G-H; mean ± SD are shown; *p* by one-way ANOVA with Tukey’s multiple comparisons). (B) Representative image of *A. baumannii* 17978 WT colonization at 10 DPI in the colon, visualized by MiPACT-HCR. Scale bar is 50 μm. blue, DAPI; magenta, WGA-lectin; green, anti-*Acinetobacter* HCR. From mice in Figure S3C. (C) Experimental design with germ-free mice. (D) *A. baumannii* 17978 CFU from fecal samples at 1, 4, and 9 DPI (n = 6 female and n = 4 male Swiss Webster mice, *p* by two-way ANOVA with Sidak’s multiple comparisons; experiment was repeated three times with similar results). (E) L-Ornithine was quantified in chow and fecal samples from untreated, post-abx, and germ-free at 0 and 10 DPI of *A. baumannii* 17978 (fecal samples are n = 10 mice shown in Figure 1G-H and Figure 3D (LOD = 5 nmol/g), chow samples are n = 3 (LOD = 1 nmol/g), mean ± SD, *p* by two-way ANOVA with Fisher’s LSD multiple comparisons only on fecal samples). (F) Principal component analysis of amino acids in fecal samples from untreated, post-abx and germ-free Swiss Webster mice at 0 and 10 DPI of *A. baumannii* 17978 with chow samples shown in each plot (n = 10 mice shown in Figure 1G-H and Figure 3D, chow n = 3). DPI, days post inoculation; MiPACT-HCR, microbial identification after passive clarity technique via hybridization chain reaction; WGA, wheat germ agglutinin; LOD, average limit of detection; PC, principal component.

To determine whether *A. baumannii* is spatially co-localized with the microbiota, *A. baumannii* was visualized *in situ* by microbial identification after passive CLARITY technique with hybridization chain reaction (MiPACT-HCR).^53,54^ Post-abx mice were mono-inoculated with either *A. baumannii* 17978 WT or Δ*astO*. There was significantly higher abundance of WT than Δ*astO* at 9 DPI (Figure S3C), confirming *A. baumannii* ornithine catabolism conferred a fitness advantage in this mono-inoculation experiment. At 10 DPI, the colon of mice inoculated with *A. baumannii* 17978 WT was harvested for MiPACT-HCR. This imaging revealed that *A. baumannii* was localized to the lumen of the colon (Figure 3B). The specificity of the *A. baumannii* HCR probe was confirmed by imaging cultures of *A. baumannii* 17978 or *E. coli* BW25113 (Figure S3D). These data support a model in which *A. baumannii* competition with the resident gut microbiota is critical for gut colonization and suggests other mechanisms, such as adherence to the host epithelium, are unlikely to contribute to *A. baumannii* gut colonization.

Collectively, the data thus far are consistent with a model in which gentamicin treatment alters the gut microbiota and disrupts initial colonization resistance to *A. baumannii*. Then as the microbiota recovers, *A. baumannii* must utilize AstO-dependent ornithine catabolism to compete with the microbiota.

Next, we sought to directly test whether *A. baumannii* requires AstO-dependent ornithine catabolism in the absence of the microbiota. Germ-free mice were orogastrically inoculated with a 1:1 mixture of the *A. baumannii* 17978 WT and Δ*astO* strains (Figure 3C). In the absence of microbiota, *A. baumannii* colonized to high levels, consistent with the idea that the intact resident microbiota confers colonization resistance (Figure 3D). There were no significant differences in colonization by WT and Δ*astO* in germ-free mice throughout the 10-day experiment (Figure 3D and S3E). Thus, AstO-dependent ornithine catabolism is dispensable in germ-free mice.

Ornithine in the gut can be produced from arginine by the host or microbiota.^44,55,56^ To determine whether the microbiota was required to produce ornithine, the amino acid metabolome was measured from the chow and feces of conventional mice (from Figure 1G-H) and germ-free mice (from Figure 3D) at 0 DPI and 10 DPI by targeted amino acid metabolomics. Ornithine levels in the chow were below the LOD of 1 nmol/g (Figure 3E); chromatograms were manually inspected to confirm that no ornithine was detected. Thus, the chow was not a major source of ornithine. In fecal samples, the levels of ornithine were variable and did not correlate with *A. baumannii* abundance or when *A. baumannii* AstO-dependent ornithine catabolism was important for colonization (Figure 3E). Germ-free mice had detectable ornithine in the fecal samples, demonstrating that the host produced ornithine in the absence of a microbiota (Figure 3E). Together, the ornithine abundance in the fecal samples did not easily explain when AstO- dependent ornithine catabolism conferred a competitive advantage to *A. baumannii*. By contrast, the amino acid metabolome of post-abx mice and germ-free mice clustered together and distinctly separate from that of untreated mice at 0 DPI (Figure 3F). This suggests that the amino acid metabolome differs between groups with varying gut microbiota and correlates to initial colonization resistance. At 10 DPI after the resident microbiota community recovered (Figure 3A, S3A, and S3B), the amino acid metabolome of post-abx mice and untreated mice cluster together (Figure 3F). Thus, the amino acid metabolome correlates with gut microbiota community recovery and requirement for *A. baumannii* ornithine utilization. Together, this suggests that overall amino acid availability may determine when *A. baumannii* depends on AstO-dependent ornithine catabolism in the gut.

### Glutamate is a preferred carbon source for *A. baumannii* that promotes gut colonization by the Δ*astO* mutant

We predicted that *A. baumannii* utilizes the ornithine niche when the resident microbiota competes for other carbon sources. *A. baumannii* carbon preferences are not described.^36^ To determine whether ornithine is a preferred carbon source for *A. baumannii, A. baumannii* 17978 was assayed in diauxic growth curves in which the non-preferred carbon source is provided in abundance with limiting concentrations of the preferred carbon source; depletion of the preferred carbon source induces a characteristic delay in growth.^57^ *Acinetobacter* species in general do not use sugars and instead use a wide variety of organic acids and amino acids as carbon sources.^36,58^ Limiting concentrations of ornithine did not induce a delay in *A. baumannii* 17978 growth with abundant glutamate or succinate, suggesting ornithine is not a preferred carbon source (Figure S4A-B). By contrast, limiting concentrations of other carbon sources (glutamate, glutamine, histidine, asparagine, or succinate) induced a delay in growth when added to excess ornithine or arginine (Figure 4A and S4C-G), showing that ornithine and arginine are not preferred carbon sources for *A. baumannii*. Further analysis showed that glutamate, glutamine, histidine, or asparagine are preferred over succinate (Figure S4H-K). In addition, the recent MDR clinical isolates *A. baumannii* AB5075 and ACICU similarly prefer amino acids such as glutamate over ornithine (Figure 4B-C), demonstrating conservation of this preference. Together, these data show *A. baumannii* prefers amino acids such as glutamate, glutamine, histidine, and asparagine over succinate, and prefers succinate over arginine and ornithine.

**Figure 4.**
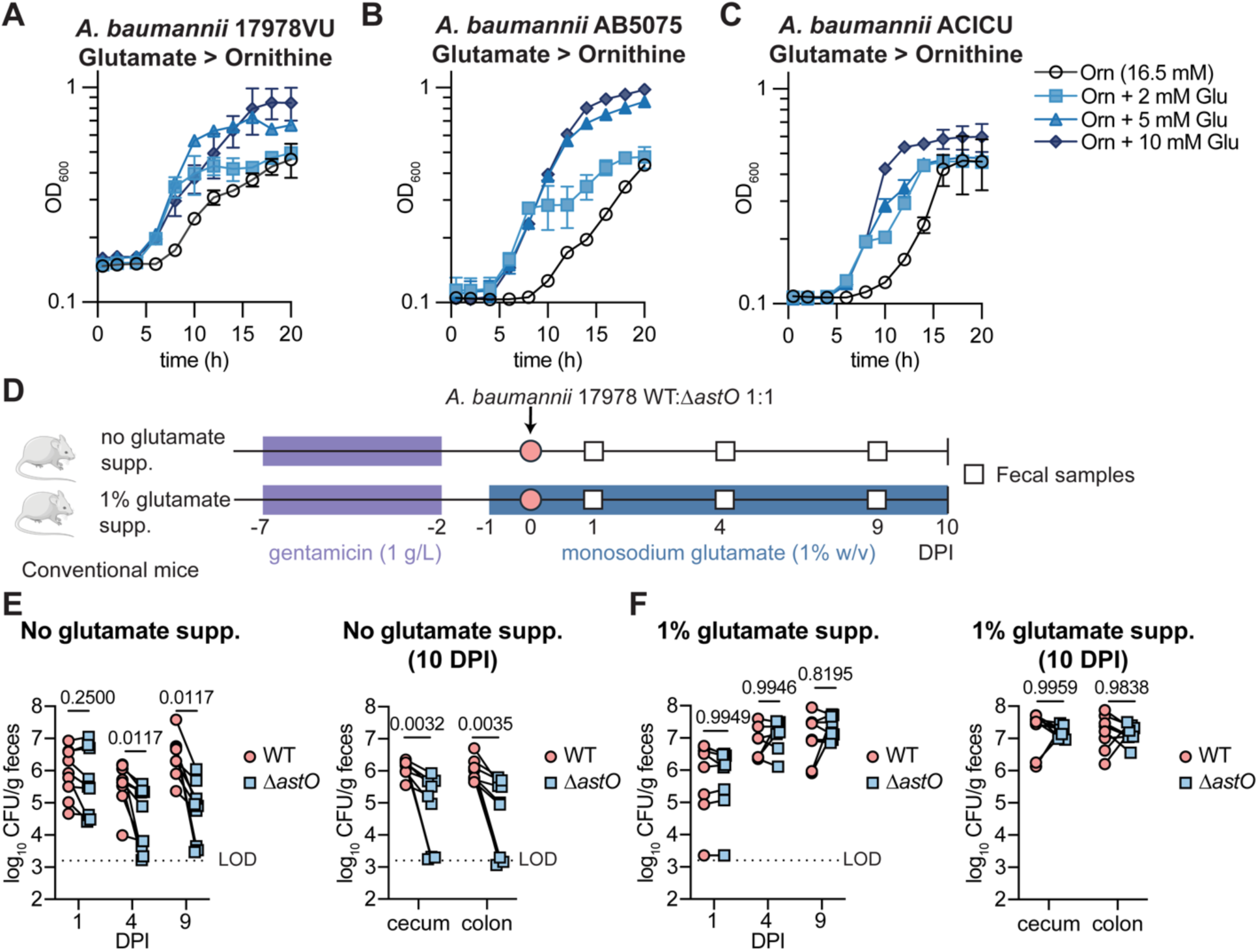
Glutamate is a preferred carbon source for *A. baumannii* that promotes gut colonization by the Δ*astO* mutant. (A-C) *A. baumannii* strains were grown in M9 media with ornithine at 16.5 mM and glutamate at indicated concentrations as the sole carbon sources. Growth was monitored by OD_600_ (n = 3; mean ± SD). (D) Experimental design for monosodium glutamate supplementation. (E-F) CFU from fecal samples at 1, 4, and 9 DPI and in the cecum and colon at 10 DPI (n = 8-9 female Swiss Webster mice, data combined from 2 independent experiments, *p* by two-way ANOVA with Sidak’s multiple comparisons). DPI, days post inoculation; LOD, average limit of detection.

We next hypothesized that dietary supplementation of a preferred carbon source such as glutamate would promote *A. baumannii* colonization without requiring AstO-dependent ornithine catabolism. To test this, monosodium glutamate (MSG), a common component of food, was supplemented in the drinking water (1% w/v) starting 1 day before inoculation and maintained throughout the 10-day experiment (Figure 4D). Consistent with previous experiments, the Δ*astO* mutant had significantly lower fecal shedding than WT at 4 and 9 DPI and in the cecum and colon at 10 DPI in mice given standard water (Figure 4E). By contrast, mice supplemented with glutamate had higher overall *A. baumannii* burdens and there was no defect for the Δ*astO* strain compared to WT (Figure 4F). Taken together, these data suggest that *A. baumannii* utilizes preferred carbon sources in the gut when available, such as early colonization in a post-abx model or with glutamate supplementation, but *A. baumannii* catabolizes ornithine for persistent gut colonization when the microbiota competes away preferred carbon sources. Furthermore, these findings suggest that dietary amino acids and protein may promote *A. baumannii* colonization in the gut.

### *A. baumannii* abundance in fecal samples from healthy human infants is increased with formula feeding

Thus far, the data presented here suggest that *A. baumannii* can colonize a low diversity microbiota such as post-abx mice and that supplementation of dietary amino acids promotes *A. baumannii* colonization. To test whether similar factors are associated in humans, we investigated *A. baumannii* abundance in fecal samples from healthy human infants. As mentioned above, *A. baumannii* can colonize the gut in neonates and has been associated with an increased risk of bloodstream infection.^17,25–27^ Prior to weaning at around 6 months of age, human infants have relatively similar diets of breastmilk or formula and have a lower complexity microbiome.^59,60^ Thus, the human infant gut represents a clinically relevant niche to examine effects of microbiota diversity and dietary protein on *A. baumannii* abundance. Shotgun metagenomic sequencing was analyzed from longitudinal fecal samples from 1-month to 24-month-old healthy human infants without exposure to antibiotics. The relative abundance of *A. baumannii* was higher in 1-month and 4-month-old infants and decreased from 4 months to 24 months (Figure 5A).

**Figure 5.**
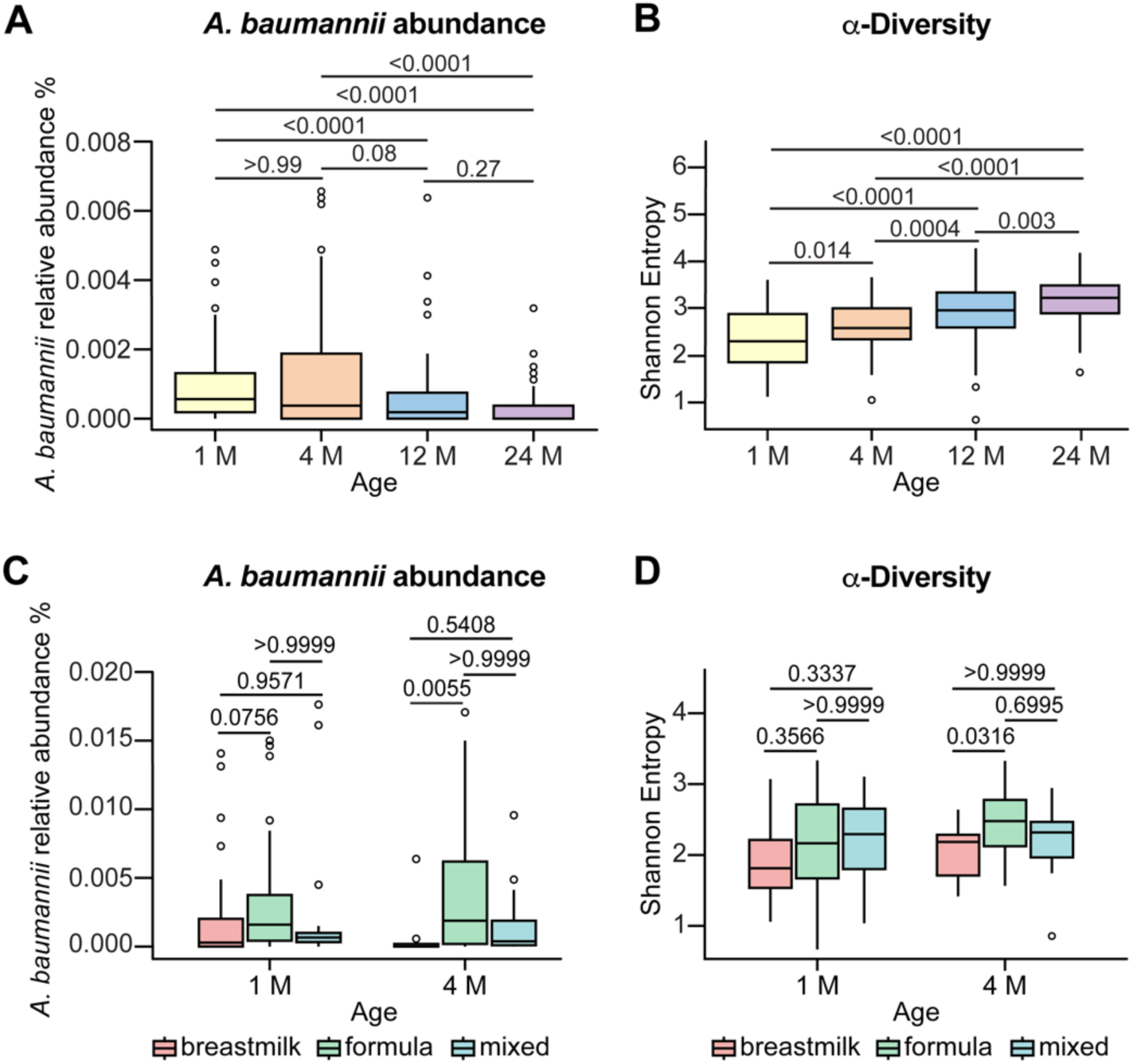
*A. baumannii* abundance in fecal samples from healthy human infants is increased with formula feeding. (A-B) Relative abundance of *A. baumannii* and microbiota α-diversity in the gut of infants age 1 month (M), 4 M, 12 M, 24 M by shotgun metagenomic sequencing (1 M, n = 104; 4 M, n = 98; 12 M, n = 104; 24 M n = 88; *p* by Kruskal-Wallis with Dunn’s post hoc test). (C-D) Relative abundance of *A. baumannii* and microbiota α-diversity in the gut of infants across feeding types at 1 M and 4 M (1 M formula, n = 45; 1 M breastmilk, n = 30; 1 M mixed, n = 29; 4 M formula, n = 62; 4 M breastmilk, n = 17; 4 M mixed, n = 19; *p* by Kruskal-Wallis with Dunn’s post hoc test). Box plots are: center line, median; box limits, upper and lower quartiles; whiskers, 1.5X interquartile range; points, outliers.

Moreover, the α-diversity of the microbiota increased with age (Figure 5B). This is consistent with the idea that as the microbiota increases in complexity, colonization resistance to *A. baumannii* develops. We next analyzed the pre-weaning infant samples by feeding type. The relative abundance of *A. baumannii* was higher with formula feeding compared to breastmilk feeding or mixed feeding in 1-month and 4- month-old infants (Figure 5C). Notably, the formula-fed infants had similar or higher α-diversity than breastmilk-fed or mixed feeding type infants (Figure 5D). Thus, the association of formula-feeding with *A. baumannii* abundance is not mediated by reduced diversity of the microbiota. Infant formula has higher protein levels than breastmilk to protect infants from essential amino acid deficiencies.^61,62^ Thus, these data support a model in which *A. baumannii* can colonize reduced diversity microbiota and that dietary protein may promote *A. baumannii* gut colonization in humans.

## Discussion

There is a growing appreciation for gut colonization as a reservoir for healthcare-associated pathogens, including *A. baumannii*. Prospective clinical studies have shown that *A. baumannii* gut colonization is a major risk factor for infection and transmission to other patients.^9,21,22^ Here, we report that ornithine catabolism is encoded by pathogenic *Acinetobacter* species and is required for *A. baumannii* gut colonization in a post-abx mouse model. *A. baumannii* AstO-dependent ornithine catabolism is dispensable in germ-free mice that lack a microbiota. Collectively, the data presented here support a model in which *A. baumannii* utilizes ornithine due to competition from the resident microbiota for preferred amino acid carbon sources such as glutamate. Furthermore, data from mice and humans suggest that dietary amino acids or protein may promote *A. baumannii* gut colonization.

Little is known about the mechanisms that *A. baumannii* uses to colonize the gut. A previous study identified a role for *A. baumannii* thioredoxin A and host secretory IgA in a short-term colonization model without antibiotics.^31^ By contrast, our study used a post-abx model and observed *A. baumannii* AstO-dependent ornithine catabolism was important for longer-term colonization as the microbiota recovered. Future work could determine whether secretory IgA also promotes long-term colonization.

Similar to our findings reported here, other studies reported that *A. baumannii* persistently colonized the mouse gut if antibiotic treatment caused microbiota disruption.^28,30^ Additionally, one study showed that 5- fluorouracil, a chemotherapy agent, promoted MDR *A. baumannii* dissemination and mortality, and these effects could be diminished by treatment with probiotic *Bifidobacterium breve*.^30^ Our data show that ornithine supplementation also promotes dissemination of MDR strain *A. baumannii* AB5075. Together, these findings suggest that environmental factors may promote dissemination of MDR *A. baumannii* from the gut to cause invasive infection.

Nutrient competition, including carbon source competition, is an important mechanism by which the gut microbiota limits pathogen colonization.^32–34,63,64^ Amino acid metabolism has been shown to be important for multiple pathogens in the gut and microbiota effects on the host.^44,65–73^ Ornithine specifically is reported to promote non-inflammatory *C. difficile* gut colonization, toxin titers, and spore formation.^45–47^ Data shown here suggests that *A. baumannii* uses ornithine in the gut when the diverse resident microbiota competes away preferred carbon sources. One recent study found that bacteria from the phylum *Bacillota* (formerly *Firmicutes*) are more likely to prefer protein as a carbon source, suggesting potential mediators of colonization resistance to *A. baumannii*.^74^ In our mouse model, multiple *Bacillota* ASVs (*e.g. Clostridia* and *Baccilli*) were depleted at 0 DPI post-abx and recovered by 9 DPI (Figure S3B), suggesting they may play a role in competitively excluding *A. baumannii* from preferred carbon sources.

Dietary amino acids and protein could play an important role in *A. baumannii* colonization and pathogen colonization in general. Ornithine and glutamate are sold commercially as post-workout supplements and MSG is a common food additive. For infants and neonates, formula typically has higher concentrations of protein than breastmilk,^62^ potentially providing more amino acid nutrients for *A. baumannii* to colonize. Previous studies showed that a high-protein diet fed to conventionally raised mice enhanced *Citrobacter rodentium* colonization^65^ and that low dietary protein protects from *C. difficile* colonization.^69^ Together with previous studies, the findings we report here are consistent with a role for amino acids and dietary protein in promoting pathogen invasion of the gut microbiota.

In conclusion, our study uncovers a metabolic strategy that *A. baumannii* uses to persist in the gut in the face of competition for nutrients by resident microbiota. The data presented here support a model in which a diverse microbiota outcompetes *A. baumannii* for preferred carbon sources, thus *A. baumannii* catabolizes ornithine to persist in an available nutrient niche. Future research into *A. baumannii* nutritional requirements in the gut and the key impacts of other members of the microbiota will provide a foundation to develop new strategies to decolonize *A. baumannii* and prevent infections. These findings have significant clinical relevance, as gut colonization is thought to be important AMR reservoir in healthcare facilities.

## Acknowledgments

We thank members of the Palmer laboratory and Erin Green (University of Chicago) for critical reading of the manuscript. We thank Stefan Green and Ashley Wu and the Genomics and Microbiome Core Facility (GMCF) at Rush University for 16S sequencing. Amino acid analysis was performed at the Microbial Culture & Metabolomics Core of the PennCHOP Microbiome Program and the Host-Microbial Repository and Analytic Core in the Center for Molecular Studies in Digestive and Liver Diseases (NIH P30 DK050306). This work was supported by startup funds from the University of Illinois Chicago to L.D.P., the Center for Microbial Medicine at the Children’s Hospital of Philadelphia to J.P.Z., an unrestricted donation from the American Beverage Foundation for a Healthy America to the Children’s Hospital of Philadelphia to support the Healthy Weight Program, the Penn Center for Nutritional Science and Medicine to G.D.W., and National Institutes of Health (NIH) National Center for Research Resources Clinical and Translational Science Program (grant no. UL1TR001878) to B.S.Z. and G.D.W. and Awards R00HL143440 to L.D.P, R01AI175223 and R01AI143641 to J.B., R35GM138369 to J.P.Z., R01AI170607 to W.H.D., R01DK107565 to B.S.Z. and G.D.W., and P30DK050306 to G.D.W.

## Author contributions

Conceptualization, X.R. and L.D.P., Methodology, X.R., R.M.C., J.D.W., C.R.T., W.H.D., and L.D.P., Investigation, X.R., R.M.C., D.B.A., E.N.V., C.R.T., J.H.G, J.D.W., K.J., O.A.T., E.S.F., and J.D.W. Writing—Original Draft, X.R. and R.M.C.; Writing—Review & Editing, X.R., R.M.C., D.A.B., E.N.V., C.R.T., J.H.G, J.D.W., K.J., O.A.T., E.S.F., B.S.Z., G.D.W., J.P.Z., W.H.D., J.B., and L.D.P.; Funding Acquisition, B.S.Z., G.D.W., J.P.Z., W.H.D., J.B., and L.D.P.

## Methods

### Resource availability

#### Lead contact

Further information and requests for resources and reagents should be directed to and will be fulfilled by the lead contact, Lauren D. Palmer.

#### Materials availability

All materials generated in this study are available from the lead contact with appropriate material transfer agreement.

#### Data and code availability

Metagenomic sequencing from healthy human infants is available in the NCBI sequence read archive (SRA) with accessions PRJNA1145027, PRJNA1106565, PRJNA1042647, and PRJNA1173239; whole-genome sequencing data for *A. baumannii* 17978VU Δ*astO*::kan is available with accession PRJNA1173193. Code used is available at https://github.com/JonWinkelman/genome_deduplication and https://github.com/JonWinkelman/dash_app_Acinetobacters/tree/main. Any additional information required to reanalyze the data reported in this paper is available from the lead contact.

### EXPERIMENTAL MODEL AND SUBJECT DETAILS

#### Ethics Statement

All animal experiments have been performed in agreement with NIH guidelines, the Animal Welfare Act, and US federal law. All animal experiments were approved by the Institutional Animal Care and Use Committee in protocol 20-165, 23-119 and 22-192 at University of Illinois at Chicago. All mice were euthanized by methods consistent with American Veterinary Medical Association (AVMA) guidelines.

#### Mouse Strains and Husbandry

##### Conventional SPF mice

Five-to-six-week-old female or male Swiss Webster mice were purchased from Charles River Lab. Swiss Webster mice were used for all 10-day experiments with *A. baumannii* 17978 inoculation gut colonization experiments and 10-week experiments with *A. baumannii* AB5075. Mice were fed autoclaved mouse diet 5L79 (PMI Nutrition International 5L79). In the 10-week experiment with *A. baumannii* 17978, six-week-old female C57BL/6J mice were purchased from The Jackson Laboratory and fed irradiated LM-485 (Teklad 7912). Ornithine in 5L79 and LM-485 was confirmed to be <1 nmol/g. For the lung infection experiment, mice were 7-week-old male C57BL/6J mice from The Jackson Laboratory fed irradiated LM-485 (Teklad 7912). For the bloodstream infection experiment, mice were 7-week-old female C57BL/6J mice from The Jackson Laboratory fed irradiated LM-485 (Teklad 7912). All mice were randomly assigned to experimental and control groups and were kept in Biologic Resources Laboratory (BRL) facility at the University of Illinois Chicago. The mice were housed with 14 h:10 h light/dark cycles, 70–76°F and 30–70% humidity.

##### Germ-free mice

Germ-free Swiss Webster (Tac:SW) WT mice were purchased from Taconic and bred in the BRL facility at the University of Illinois Chicago in a room with 14 h:10 h light/dark cycles and 70–76°F and 30–70% humidity. Mice were kept in isolators purchased from Park Bioservices LLC. Mice were fed autoclaved mouse diet 5L79 (PMI Nutrition International 5L79) and autoclaved super Q water *ad libitum* and were 12-16 weeks of age. Germ-free condition was tested at least once a month with aerobic liquid cultures (brain heart infusion, BHI), solid cultures (blood agar plates, Thermo Sci Remel, R01200), fungal cultures (Sabouraud slants), anaerobic liquid (BHI) and solid cultures (*Brucella* agar, Thermo Sci Remel, R01254) from isolators’ swabs, fecal samples, and fungal traps placed inside the isolators. Fecal samples were also tested with Gram staining and qPCR to detect bacterial DNA.

#### Bacterial strains and routine culture conditions

*A. baumannii* strains, *E. coli* K12, and *Pseudomonas aeruginosa* PAO1 were grown in LB ((10 g/L tryptone, 5 g/L yeast extract, 10 g/L sodium chloride) or LB agar (10 g/L tryptone, 5 g/L yeast extract, 10 g/L sodium chloride, 15 g/L agar) at 37°C. *A. baylyi*, *A. gyllenbergii* and *A. colistiniresistens* were grown in LB or LB agar at 30°C. Antibiotics were used at the following concentrations: carbenicillin 75 mg/L, kanamycin 40 mg/L, chloramphenicol 15 mg/L. For experiments with mice, the antibiotics used for selective plating were used at the following concentrations: carbenicillin 50 mg/L, kanamycin 40 mg/L, chloramphenicol 5 mg/L.

### METHOD DETAILS

#### Bacterial strain construction

The strains, recombinant DNA, and oligonucleotides used in this study listed in Table S1. *A. baumannii* 17978 is ATCC 17978VU.^75^ Mutants were generated by allelic exchange with sucrose counterselection using pFLP2.^76^ Approximately 1,000 bp of DNA in both the 5’ and 3’ flanking regions surrounding targeted genes was amplified using *A. baumannii* genomic DNA as a PCR template.

Oligonucleotides were purchased from Integrated DNA Technologies. The kanamycin resistance marker was amplified from the vector pKD4.^77^ The PCR products were cloned into the pFLP2 vector using HiFi Assembly (New England Biolabs). The pFLP2 constructs were then introduced into WT *A. baumannii* by electroporating and selecting for Kn^R^, resulting colonies were further screened for Carb^R^ and sucrose^S^. Then the sucrose sensitive and PCR-confirmed merodiploids were plated to LB agar containing 10% sucrose to select for loss of the plasmid. Resulting colonies were screened for Kn^R^ and Carb^S^. Generation of Δ*astO*::Kn was confirmed by multiple PCRs and whole genome sequencing. For construction of the site mutation *astA^H^*^229^*^A^*, 1,000 bp of DNA in both the 5’ and 3’ flanking regions surrounding *astA* nucleotides encoding H229A L125A was amplified by PCR. These PCR products were cloned into the pFLP2 vector using HiFi Assembly. After sequence confirmation, the resulting plasmid was introduced into WT by electroporating and selecting for Carb^R^ and screening for sucrose^S^. Then the sucrose sensitive clones were plated onto agar containing 10% sucrose to select for loss of the plasmid. The resulting colonies were screened for Carb^S^. The site mutation strain *astA^H^*^229^*^A^ ^L^*^125^*^A^* was confirmed by amplifying the region by PCR and sequencing. For construction of the site mutation *astA^H^*^229^*^A^ ^L^*^125^*^A^*Δ*astO*::Kn, the pFLP2- *astO*::Kan plasmid was introduced into *ast1^H^*^229^*^A^ ^L^*^125^*^A^* by electroporating and selected as described above. For the *astO* complementation vector, open reading frames (ORFs) were amplified by PCR and cloned into pWH1266 vector with a constitutive *A. baumannii rpsA* promoter (*rpsAp*) by Hifi Assembly. The resulting plasmid was introduced into *A. baumannii* Δ*astO* by electroporating and selected for Carb^R^ to generate the complementation strain. To construct the *A. baumannii att::*mTn*7* marked WT strain, the pKNOCK-mTn*7* was transformed into *A. baumannii* WT by four parental conjugal mating strategy as described previously.^78^

#### Bacterial growth curves

*A. baumannii* strains, *A. nosocomialis, P. aeruginosa*, and *E. coli* were grown in 3 mL LB at 37°C overnight with shaking at 180 rpm. Cultures were diluted 1/100 into PBS, then further diluted 1/100 into 99 μL carbon-free nitrogen-free M9 minimal medium with the indicated carbon source (16.5 mM) and nitrogen source (18.6 mM) in a flat bottom 96-well plate (Fisher) and covered with Breathe-Easy sealing membrane. Growth was monitored in a BioTek Synergy 2 plate reader or Epoch 2 plate reader by optical density at 600 nm (OD_600_) at 37°C with shaking. *A. baylyi*, *A. glyllenbergii*, and *A. colistiniresistens* were cultured as described but at 30°C.

#### Analysis of *ast2* operon prevalence among *A. baumannii* strains

Protein sequence identity was determined with EMBOSS needle.^79^ The prevalence of the *ast2* operon was assessed in the previously described set of deduplicated *A. baumannii* genomes.^43^ Briefly, 7,431 genome assemblies were downloaded from NCBI. Genomes with contig N50 scores in the lowest 20% were removed, reducing the dataset to 5,945 genomes. Due to the presence of many closely related genomes, a deduplication process was conducted using Average Nucleotide Identity (ANI) values, estimated via Mash 2.34.^80^ A custom Python script iterated through the genomes, comparing their Mash distances, and discarded genomes with lower N50 scores if they were too similar (based on a threshold distance of 0.006). Additionally, ten genomes identified as outliers were excluded. The code used is previously published and available at https://github.com/JonWinkelman/genome_deduplication.^43^ This resulted in 229 proteomes from filtered and deduplicated *Acinetobacter* genomes (Table S2), with four outgroup species (*A. nosocomialis, A. colistiniresistens*, *A. gyllenbergii*, and *A. baylyi*) that were used to root the species tree. Orthofinder generated hierarchical orthologous groups (HOGs) for each internal node of the species tree, representing proteins descended from a common ancestral gene. HOGs linked to the last common ancestor of all *A. baumannii* species were used. Details on analysis and visualization are available at https://github.com/JonWinkelman/dash_app_Acinetobacters/tree/main.

#### Phylogenetic trees

Trees were inferred with OrthoFinder version 2.5.5^81,82^ or RAxML.^83^ When inferring gene/protein trees, protein alignments for RAxML were produced with muscle 5.1.^84^ The RAxML command was configured to perform phylogenetic analysis using the PROTGAMMAAUTO model, which automatically selects the best-fitting protein substitution model combined with the Gamma model of rate heterogeneity. Rapid Bootstrap analysis and searches for the best-scoring Maximum Likelihood tree were performed in a single run. Random number seeds provided for both the rapid bootstrap analysis (-x 123) and the parsimony inference step (-p 256). The number of bootstrap replicates was determined automatically using the autoMRE option, which stops generating replicates once a sufficient number have been produced to yield reliable support values.

#### Mouse experiments

Purchased mice were first acclimated in the facility for one week. Conventional mice were then treated with 1 g/L gentamicin in the drinking water for 5 days (day -7 to -2); gentamicin water was replaced every 2-3 days; at day -2, all mice were given normal drinking water. For supplementary ornithine or glutamate water experiment, mice received supplementary ornithine HCl (1% w/v) or monosodium glutamate monohydrate (1% w/v) in the drinking water from day -1 before inoculation with *A. baumannii* until the end of the experiment. *A. baumannii* 17978 WT and Δ*astO* mutant strain were grown in 10 mL LB at 37°C with shaking for 16 h. *A. baumannii* AB5075 was incubated in 10 mL LB at room temperature without shaking overnight, then incubated at 37°C with shaking for 3.5 h. Subsequently, the bacteria were centrifuged at 4°C, 4000 x *g* for 7 min, washed in PBS twice, and resuspended in PBS to 1 x 10^10^ colony forming units (CFU)/mL. On day 0, mice were orally gavaged with 0.1 mL (1 x 10^9^ CFU) of the indicated strains and placed in a clean cage. Cages were changed at least weekly. For *A. baumannii* CFU enumeration, fecal samples were weighed, resuspended in 1 mL PBS, serially diluted, and plated to LB plates supplemented with appropriate antibiotics. CFU were enumerated on LB agar with 5 mg/L chloramphenicol and 50 mg/L carbenicillin (*A. baumannii* 17978 WT *att*::mTn*7)*, with 5 mg/L chloramphenicol and 40 mg/L kanamycin (*A. baumannii* 17978 Δ*astO*::Kn), or with 15 mg/L chloramphenicol (*A. baumannii* AB5075 WT). Fecal samples were collected at 0 DPI and plated to confirm no *A. baumannii* prior to inoculation. Mice were euthanized with CO_2_. Cecum and colon were collected in 1 mL PBS and the spleen was collected in 0.7 mL PBS and homogenized by Bullet Blender with stainless steel beads (NextAdvance). The serial dilutions of homogenates were plated on LB agar with appropriate antibiotics. For the germ-free mouse experiment, 10^7^ and 10^9^ *A. baumannii* inocula were tested and the data were combined.

For lung infection, *A. baumannii* 17978 WT and Δ*astO* mutant strains were grown overnight in 3 mL LB with shaking at 180 rpm at 37°C for 16 hours. The overnight cultures were then subcultured 1:100 into 10 mL LB and grown for 3.5 h. *A. baumannii* were harvested by centrifugation, washed twice in PBS, and resuspended in PBS at 1 x 10^10^ CFU/mL. Mice were anesthetized with intraperitoneal injection of ketamine/xylazine diluted in PBS. Mice were then inoculated intranasally with 1:1 ratio of *A. baumannii* WT and Δ*astO* at approximately 3 x 10^8^ CFU in 30 μL. At 1 DPI, mice were euthanized with CO_2_ and organs (lungs, spleen, liver, heart, kidneys and nasal cavity) were collected. Organs were homogenized in PBS using a NextAdvance Bullet Blender tissue homogenizer on setting 8. CFU were enumerated on LB agar (total *A. baumannii* 17978), and LB agar with 40 mg/L kanamycin (Δ*astO*::Kn); WT CFU were determined by subtraction.

For retroorbital infection, *A. baumannii* 17978 WT and Δ*astA*Δ*astO* double mutant strains were prepared as described for lung infections. Mice were anesthetized by isoflurane. Mice were then inoculated retro-orbitally with a 1:1 ratio of *A. baumannii* 17978 WT and Δ*astA*Δ*astO* at approximately 5 x 10^8^ CFU in 50 μL. At 1 DPI, mice were euthanized, organs (lungs, spleen, liver, heart, and kidneys) were collected, and CFU were enumerated as described above.

#### MiPACT-HCR imaging

Mice were mono-inoculated with either WT or Δ*astO A. baumannii* 17978 as described above. After euthanasia, the whole colon was fixed in methacrylate (10% acetic acid, 30% chloroform, and 60% methanol). The organ was then embedded, sectioned and stained as previously described.^54^ Briefly, colons were embedded twice in B4P1 (bis-acrylamide/paraformaldehyde mixture) to ensure retention of lumen contents during clearing. After clearing, the samples were stained with DAPI, and lectin (wheat germ agglutinin [WGA]) conjugated to AlexaFluor-647 (Thermo Scientific). Hybridization chain reaction (HCR) was performed with an *Acinetobacter*-specific FISH probe (Aci-16s 729)^85^ with a B4 overhang, and amplified by fluorescent DNA hairpins conjugated to Alexa Fluor-488 (Molecular Instruments).

Imaging was performed using a Zeiss LSM 780 confocal microscope or a Zeiss LSM 880 confocal microscope with a Plan-Apochromat 10×/0.45-numerical aperture M27 objective. Scanned images were processed using Imaris imaging software (Bitplane) or the FIJI distribution of ImageJ.

#### Mouse microbiota sequencing and analysis

Mouse fecal samples were collected and immediately stored at -80°C until processing. DNA was extracted using by QIAamp PowerFecal Pro DNA Kit according to the manufacturer’s instructions.

Library preparation and sequencing was conducted at the Rush University Genomics and Microbiome Core Facility (GMCF). Microbial community characterization was performed using two-stage 16S rRNA gene amplicon sequencing, similar to methods previously described.^86,87^ Briefly, gDNA was amplified with primers targeting the V1-V3 variable regions of microbial 16S rRNA genes using a mixture of 27F primers and 534R primers. Forward primers were a combination of multiple versions of the 27F primer, as described previously,^88^ and including 27F-YM, 27F-Bor, 27F-Bif, 27F-Chl, and 27F-Ato forward primer variants. All primers contained Fluidigm common sequence linkers (CS1, ACACTGACGACATGGTTCTACA, for forward primers; CS2, TACGGTAGCAGAGACTTGGTCT, for reverse primers). PCR reactions were conducted in 10 μl volumes with repliQa HiFi Tough Mix (QuantaBio). PCR conditions included an initial denaturation at 98°C for 2 minutes, followed by 24 cycles of 98°C for 10 seconds, 56°C for 2 seconds, and 68°C for 1 second. Subsequently, a second PCR reaction was performed using 1 μl of the product from the first stage as the input without purification. The second-stage primers were from the Access Array for Illumina barcoding system (Fluidigm, San Francisco, CA, USA) and contained Illumina sequencing adapters, sample-specific barcodes, and CS1 and CS2 linkers at the 3’ ends of the oligonucleotides. The PCR conditions for these second reactions were an initial denaturation at 98°C for 2 minutes, followed by 8 cycles of 98°C for 10 seconds, 60°C for 1 second, and 68°C for 1 second. After the 2nd stage amplification, samples were pooled approximately equimolarly and sequenced on an Illumina MiSeq sequencer and employing a V3 kit with paired-end 2 x3 00 base sequencing reads. Demultiplexing was performed on the sequencing instrument.

The resulting paired-end FASTQ files were merged using the Paired-End read merger (PEAR) algorithm.^89^ Merged data were then quality trimmed (≥ 20) and filtered for sequences ≥ 450 bases. The trimmed and filtered sequences were exported as FASTA and analyzed using nf-core/ampliseq version 2.1.0.^90,91^ Amplicon sequence variants (ASVs) taxonomies were assigned by DADA2 using SILVA-138 database.^92–94^ α-diversity, β-diversity, and relative abundance bar plots were generated with QIIME 2.^95^

Non-metric MultiDimensional Scaling (NMDS) plot of Jaccard β-diversity was generated with the R package vegan (version 2.6-8).

#### Amino acid metabolomics and quantification

Samples were pre-weighed and homogenized in sterile PBS and then sterilized with 0.22-µm filters. Amino acids were derivatized using the Waters AccQ-Tag Ultra Amino Acid Derivatization Kit (Waters Corporation, Milford, MA) and analyzed using the UPLC AAA H-Class Application Kit (Waters Corporation, Milford, MA) according to manufacturer’s instructions. Amino acids were quantified using a Waters Acquity uPLC System with an AccQ-Tag Ultra C18 1.7 μm, 2.1 × 100 mm column and a photodiode detector array with a standard curve. Quality control checks (blanks and standards) were run every eight samples. For ornithine quantification from mouse chow, ornithine peaks were manually inspected and confirmed to be absent in the chow samples. All chemicals and reagents used were mass spectrometry grade.

#### Shotgun metagenomics of human fecal samples

Samples were from infants who had never received antibiotics and had no history of health problems at the Children’s Hospital of Philadelphia as part of the Infant Growth and Microbiome (IGraM) Study, a prospective, longitudinal cohort study of pregnant African American women and their infants.

The IGraM study was approved by the Committee for the Protection of Human Subjects (Institutional Review Board) at Children’s Hospital of Philadelphia.^96^ The data used here are publicly available. Human DNA isolation and shotgun metagenomic sequencing were performed as previously described.^97^ Briefly, DNA was extracted from feces and negative controls using DNeasy PowerSoil HTP 96 Kit following the manufacturer’s instructions. Shotgun libraries were generated from 1 ng of DNA using the NexteraXT kit (Illumina). Libraries were sequenced on the Illumina HiSeq using 2 × 125 bp chemistry in High Output mode.

Shotgun metagenomic data were analyzed using Sunbeam v.2.0.1. The taxonomic count matrix from Kraken was subjected to quality filtering using the R package phyloseq.^98^ Taxa that were present in fewer than 25% of samples and less than 1000 cumulative reads were removed. Samples with less than 1 million reads were also removed from the dataset prior to analysis. Statistics on *A. baumannii* relative abundance between age groups and feeding type were performed using a Kruskal-Wallis test, and group comparisons were obtained using a Dunn’s post hoc test using the R package rstatix. P values displayed in figures were corrected for the false discovery rate.

Shannon entropy scores were calculated using the estimate richness function in the phyloseq package. Statistics on Shannon entropy scores between age groups were performed using a Kruskal Wallis-test, and group comparisons were obtained using a Dunn’s post hoc test using the package rstatix. P values were corrected post hoc for the false discovery rate. Graphs were generated using the R package ggplot2.

#### Statistical analysis

All statistical analyses were performed in R v.4.0.2 or Prism v.10.1.1 (GraphPad). Statistical tests used are indicated in each figure legend. Figures were prepared in Adobe Illustrator 28.1.

## Supplemental information

SI Document with Figures S1–S4, Table S1 Table S2: *A. baumannii astR*, *astN, astO, astP* frequency

