## Supplemental Information for "Amino acid competition shapes *Acinetobacter baumannii* gut carriage"

Lauren D. Palmer

835 S Wolcott Ave

MSB E703

Chicago, IL 60612

#Present address: American University of the Caribbean, Cupecoy, Sint Maarten

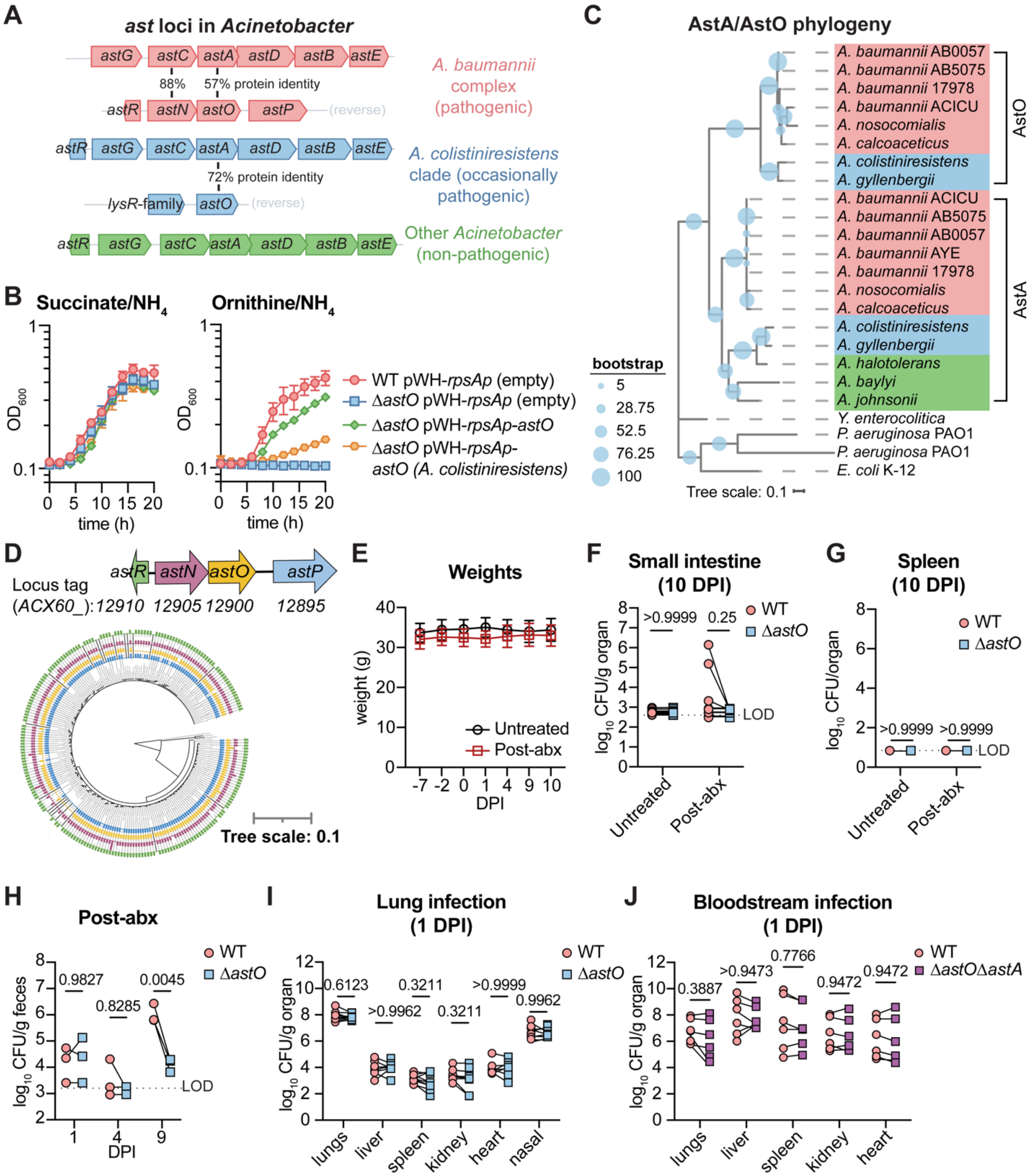


**Fig S1. Evidence of evolution at the second *ast* locus in *Acinetobacter* spp. and requirement for *A. baumannii* gut colonization.**

Corresponds with Figure 1

(A) *ast* loci in *Acinetobacter* clades. Members of the *abc* clade such as *A. baumannii* 17978 encode the second *ast* locus with *astR*, and *astNOP* on the negative strand (red). Members of the *A. colistiniresistens* clade encode the *astGCADBE* locus with *astR* divergently transcribed; the second *ast* locus has only *astO* and a divergent LysR-family regulator gene (blue). Other non-pathogenic *Acinetobacter* spp. only one *ast* locus with *astR* and *astGCABDE* (green).

(B) Growth of *A. baumannii* 17978 WT pWH (empty vector), Δ*astO* pWH, Δ*astO* pWH-*astO* (*A. baumannii*) and Δ*astO* pWH-*astO* (*A. colistiniresistens*) grown in M9 minimal media with succinate or ornithine as the sole carbon source. Growth was monitored by $OD_{600}$ measurement for 20 h (n = 3, mean +/- SD).

(C) Phylogenetic tree of AstA and AstO proteins (tree scale in amino acid substitutions).

(D) Second *ast* locus genes and their corresponding copy numbers mapped to an *A. baumannii* species phylogenetic tree generated from 233 de-duplicated published *A. baumannii* and *Acinetobacter* genomes (see Table S2; tree scale in amino acid substitutions).

(E) Weight of female Swiss Webster mice in Figure 1G-H (n = 10, mice combined from 2 independent experiments, mean ± SD).

(F) *A. baumannii* 17978 CFU in the small intestine at 10 DPI from mice shown in Figure 1G-H (n = 10, mice combined from 2 independent experiments, *p* by Wilcoxon test with Holm-Sidak's multiple comparisons).

(G) *A. baumannii* 17978 CFU in the spleen at 10 DPI from female Swiss Webster mice shown in Figure 1G-H (n = 10, mice combined from 2 independent experiments; *p* by Wilcoxon test with Holm-Sidak's multiple comparisons).

(H) Male Swiss Webster mice were administered gentamicin as in Figure 1F and at 0 DPI were inoculated with 1:1 *A. baumannii* 17978 WT and Δ*astO*. CFU were enumerated from feces at 1, 4, and 9 DPI (n = 3, *p* by two-way ANOVA with Sidak's multiple comparisons).

(I) Female C57BL/6 mice were intranasally inoculated with 1:1 *A. baumannii* 17978 WT and Δ*astO*. CFU were enumerated at 1 day post inoculation (DPI) (n = 10, *p* by Wilcoxon test with Holm-Sidak's multiple comparisons).

(J) Male C57BL/6 mice were retroorbitally inoculated with 1:1 *A. baumannii* 17978 WT and Δ*astA*Δ*astO*. CFU were enumerated at 1 DPI (n = 6, *p* by Wilcoxon test with Holm-Sidak's multiple comparisons).

DPI, days post inoculation; LOD, average limit of detection.

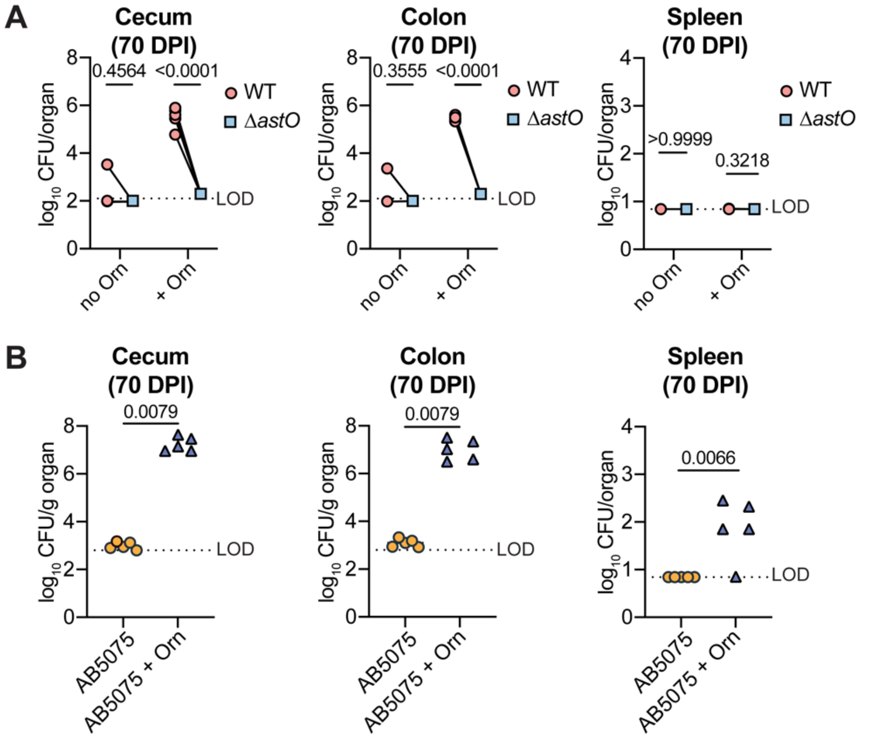


**Figure S2 Supplemental ornithine promotes long-term *A. bauma*nnii gut colonization**

Corresponds with Figure 2

(A) Female C57BL/6J mice (from Figure 2B) were euthanized at 70 DPI and *A. baumannii* CFU were enumerated from the cecum, colon and spleen (n = 5, *p* by two-way ANOVA with Sidak's multiple comparisons).

(B) Female Swiss Webster mice (from Figure 2D) were euthanized at 70 DPI and *A. baumannii* 5075 CFU were enumerated from the cecum, colon and spleen (n = 5, *p* by two-way ANOVA with Sidak's multiple comparisons).

DPI, days post infection; LOD, average limit of detection.

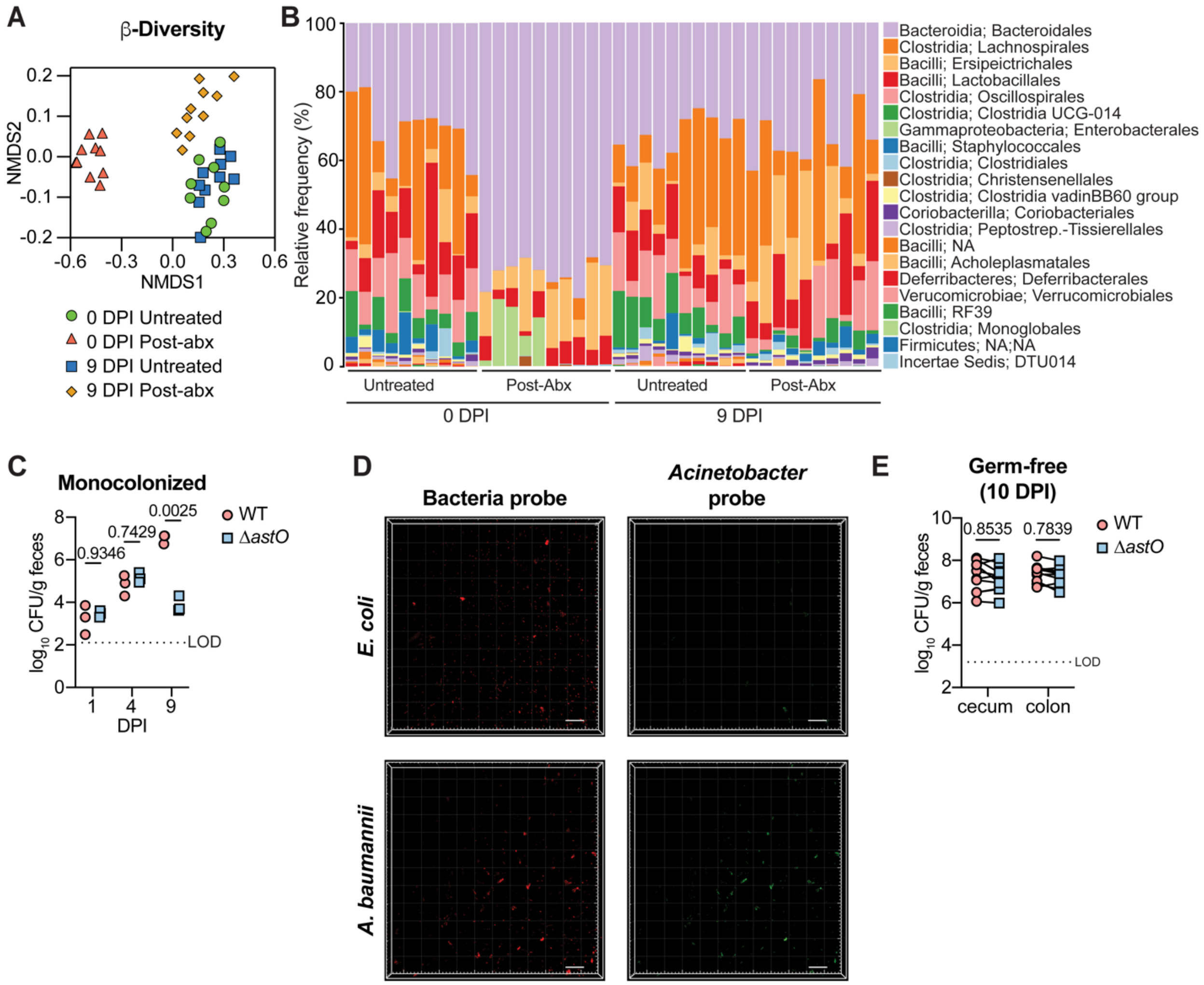


**Figure S3 WT *A. baumannii* outcompetes Δ*astO* by 9 DPI.**

Corresponds with Figure 3

(A) β-diversity NMDS plot of 16S rRNA gene profiling at 0 and 9 DPI in the feces of untreated and post-abx female Swiss Webster mice inoculated with *A. baumannii* 17978 WT and Δ*astO* and shown in Figure 1G-H (n = 10).

(B) Relative abundance of bacterial ASV identified by 16S rRNA gene sequencing at 0 and 9 DPI in the feces of untreated and post-abx female Swiss Webster mice shown in Figure 1G-H (n = 10).

(C) Mice were mono-inoculated with *A. baumannii* 17978 WT or Δ*astO*. CFU were enumerated from fecal samples at 1, 4 and 9 DPI (n = 3 female Swiss Webster mice, *p* by two-way ANOVA with Sidak's multiple comparisons). Mice inoculated with WT were used for MiPACT-HCR imaging shown in Figure 4B.

(D) MiPACT-HCR imaging of bacterial cultures to assess specificity of anti-*Acinetobacter* probe Aci16s 729. Scale bar is 50 µm; green, anti-*Acinetobacter* HCR probe; red, general bacterial HCR probe eub338.

(E) GF mice were euthanized at 10 DPI and CFU were enumerated (n = 10, *p* by two-way ANOVA with Sidak's multiple comparisons).

NMDS, non-metric multidimensional scaling; DPI, days post infection; ASV, amplicon sequence variants; LOD, limit of detection; MiPACT-HCR, microbial identification after passive clarity technique via hybridization chain reaction.

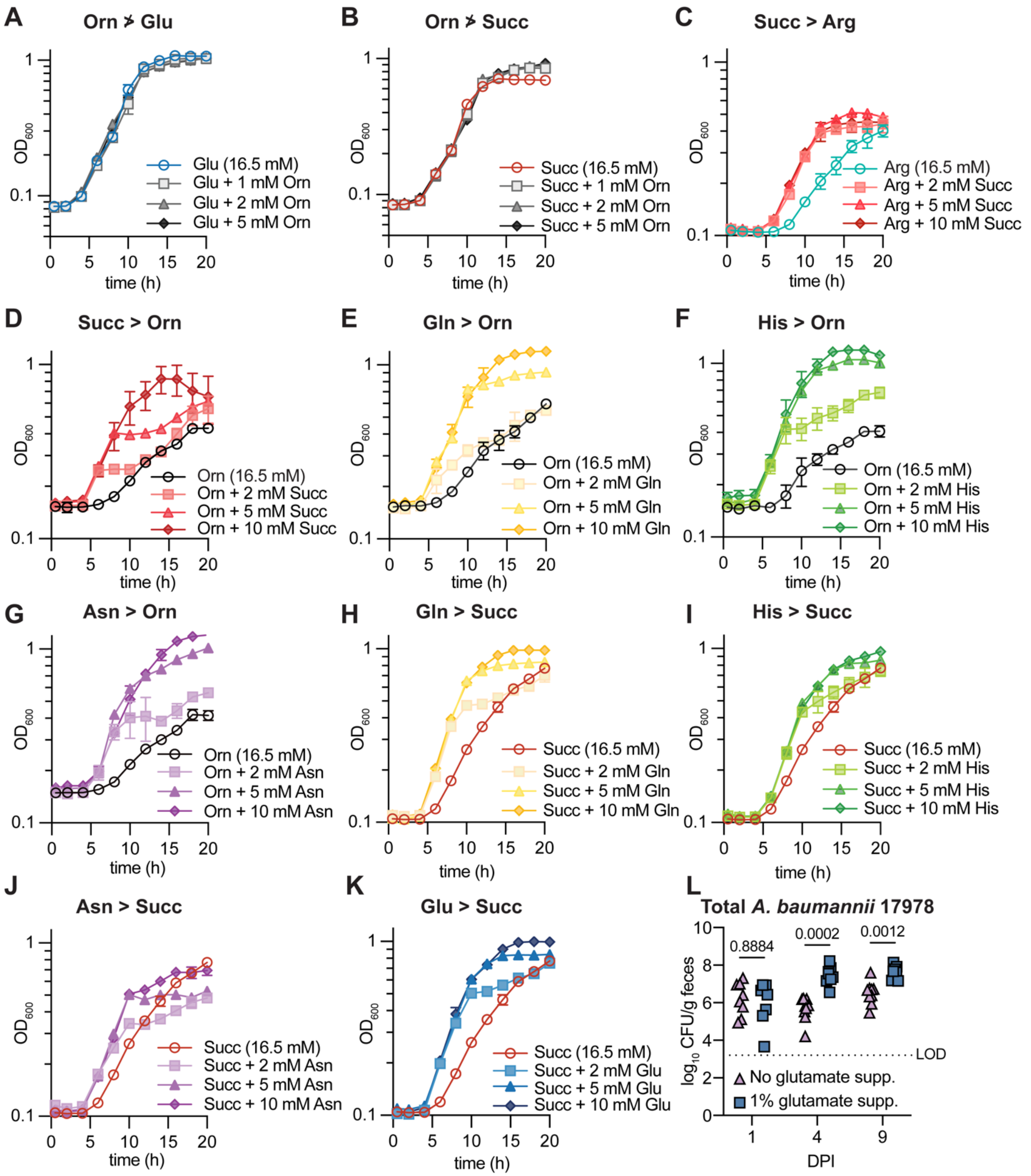


**Figure S4** ***A. baumannii*** **preferred carbon source**

Corresponds with Figure 4

(A-K) *A. baumannii* 17978 WT growth curve in M9 media containing 16.5 mM carbon source and indicated additions as the sole carbon sources. Growth was monitored by $OD_{600}$ (n = 3, mean ± SD). *A. baumannii* preferred carbon source is Glu, Gln, Asn, His > Succ > Arg, Orn

(L) Post-abx mice were co-inoculated with *A. baumannii* 17978 WT and Δ*astO* and one group was supplemented with 1% monosodium glutamate in the drinking water. Total *A. baumannii* CFU were enumerated from fecal samples at 1, 4 and 9 DPI (n = 8-9 female Swiss Webster mice also shown in Figure 4E-F, $p$ by two-way ANOVA with Sidak's multiple comparisons). Orn, ornithine; Glu, glutamate; Succ, succinate; Gln, glutamine; His, histidine; Asn, asparagine; Arg, arginine. DPI, days post infection; LOD, limit of detection.

**Table S1.** Strains, plasmids, and oligonucleotides.

| REAGENT or RESOURCE | SOURCE | IDENTIFIER |
|---|---|---|
| Bacterial and virus strains | | |
| *Acinetobacter baumannii* ATCC 17978VU | ATCC (Wijers et al., 2021) | LP486 |
| *Acinetobacter baumannii* ATCC 17978VU*::mTn7* | This study | LP526 |
| *A. baumannii* Δ*astO* deletion strain used in IN infection (old one) | This study | LP720 |
| *A. baumannii* Δ*astO* Δ*astA* deletion strain used in RO infection | This study | LP516 |
| *Acinetobacter baumannii* ATCC 17978VU Δ*astO::kan* | This study | LP814 |
| *Acinetobacter baumannii* ATCC 17978VU *astA1* $^{L125A\ H229A}$ | This study | LP838 |
| *Acinetobacter baumannii* ATCC 17978VU *astA1* $^{L125A\ H229A}$Δ*astO::kan* | This study | LP872 |
| *Acinetobacter baumannii* ATCC 17978VU pWH1266-$P_{rpsA}$ | This study | LP731 |
| *Acinetobacter baumannii* ATCC 17978VU Δ*astO::kan* pWH1266-$P_{rpsA}$ | This study | LP732 |
| *Acinetobacter baumannii* ATCC 17978VU Δ*astO::kan* pWH1266-$P_{rpsA}$-*astO* | This study | LP1019 |
| *Acinetobacter baumannii* ATCC 17978VU Δ*astO::kan* pWH1266-$P_{rpsA}$-*astO(A. colistiniresistens)* | This study | LP1020 |
| *Acinetobacter baumannii* ATCC AYE | ATCC | LP13 |
| *Acinetobacter baumannii* 0057 | Robert Bonomo (Case Western Reserve University) | LP14 |
| *Acinetobacter baumannii* ACICU | M. Stephen Trent (University of Georgia) | LP293 |
| *Acinetobacter baumannii* ABUW AB5075 | Colin Manoil (University of Washington) | LP345 |
| *Acinetobacter nosocomialis* | Mario Feldman (Washington University at St. Louis) | LP116 |
| *Acinetobacter baylyi* ADP1 | ATCC | LP459 |
| *Acinetobacter colistiniresistens* | DSMZ | LP697 |
| *Acinetobacter gyllenbergii* | DSMZ | LP698 |
| *Escherichia coli* K12 | Maria Hadjifrangiskou (Vanderbilt University Medical Center) | LP235 |
| *Escherichia coli* BW25113 | Matthew Chapman (University of Michigan) | BW25113 |
| *Pseudomonas aeruginosa* | Andrea Battistoni (University of Rome Tor Vergata) | LP346 |
| Oligonucleotides | | |
| gttaaaaaggatcgatcctctagaggatcCTATACAAATGAACCCGTTCTAC | | *astO*_up_F |
| agctccagcctacacGCTGCTTGTTCATCCTTTTG | | *astO*_up_R |
| gaggatattcatatgGCGGAAATTACTATACATTTCAC | | *astO*_dn_F |
| tgaccatgattacgaattcgagctcggtacCATTTGGAGAAGTAAACCC | | *astO*_dn_R |
| GTGTAGGCTGGAGCTGCTTC | | pKD4-FRTfrag_F |
| CATATGAATATCCTCCTTAGTTCCTATTC | | pKD4-FRTfrag_R |
| GTTAAAAAGGATCGATCCTCTAGAatgtacttacgactgcaaaag | | *astA1*_up_F |
| aaTGCtgtacagagctcactac | | *astA1*_L125A_R |
| agtgagctctgtacaGCAttttta | | *astA1*_L125A_F |
| tgtggTGCcatttttccaatca | | *astA1*_H229A_R |
| aaaaatgGCAccacatactttgcc | | *astA1*_H229A_F |
| AATTCGAGCTCGGTACCtacaactgtgttaccagcaagt | | *astA1*_dn_F |
| TTATCAGGTATATCctcgagatgatgattattcgttacattgaac | | Com_*astO*_F |
| GGGCATCGGTCGACggtaccattaacagccattcgaaaat | | Com_*astO*_R |
| TTATCAGGTATATCctcgagATGATGCTGATTCGTTATATCA | | Com_*astO(A.colistiniresistens)*_F |
| GGGCATCGGTCGACggtaccTCAATTTTTCTTTGGAATATTG | | Com_*astO(A.colistiniresistens)*_R |

| | | |
|---|---|---|
| gttaaaaaggatcgatcctctagaggatcCTATACAAATGAACCCGTTCTAC | | *astO*_up_F |
| agctccagcctacacGCTGCTTGTTCATCCTTTTG | | *astO*_up_R |
| gaggatattcatatgGCGGAAATTACTATACATTTCAC | | *astO*_dn_F |
| tgaccatgattacgaattcgagctcggtacCATTTGGAGAAGTAAACCC | | *astO*_dn_R |
| GTGTAGGCTGGAGCTGCTTC | | pKD4-FRTfrag_F |
| CATATGAATATCCTCCTTAGTTCCTATTC | | pKD4-FRTfrag_R |
| AGAGTTTGATYMTGGCTCAG | | 16s-CS1_27F-YM |
| AGAATTTGATCTTGGTTCAG | | 16s-CS1_27F-Chl |
| AGAGTTTGATCCTGGCTTAG | | 16s-CS1_27F-Bor |
| AGGGTTCGATTCTGGCTCAG | | 16-CS1_27F-Bif |
| AGAGTTCGATCCTGGCTCAG | | 16s- CS1_27F-Ato |
| ATTACCGCGGC GCTGG | | 534R |
| **Recombinant DNA** | | |
| pFLP2 | Hoang et al., 1998 | N/A |
| pKD4 | Datsenko and Wanner, 2000 | N/A |
| pKNOCK-mTn7-Amp | Carruthers *et al.*, 2013 | N/A |
| pWH1266 | Hunger *et al.*, 1990 | N/A |
| pWH1266-*rpsAp* | Palmer *et al.*, 2020 | pLDP29 |
| pWH1266-*rpsAp-astO* | This study | pLDP171 |
| pWH1266-*rpsAp-astO (colistiniresistens)* | This study | pLDP181 |
| pFLP2-Δ*astO::Kn* | This study | pLDP94 |
| pFLP2-*astA1*$^{L125A\ H229A}$ | This study | pLDP216 |
